# Coupled Modeling of Tumor Volume and ctDNA Dynamics in HPV-Associated Anal Squamous Cell Carcinoma

**DOI:** 10.64898/2026.11.01.763528

**Authors:** Phebe M. Havor, James Jabalee, Brandon M. Huffman, James M. Cleary, Renee Brady-Nicholls

## Abstract

Response to anti-PD-1 immunotherapy in advanced HPV-associated anal squamous cell carcinoma (ASCC) varies and is assessed by infrequent imaging that lags behind tumor biology. Circulating tumor DNA (ctDNA) can be sampled frequently and reflects near-real-time tumor activity yet is rarely built into mechanistic models of treatment response. Using longitudinal tumor volume and HPV ctDNA from 26 patients treated with pembrolizumab, we found that baseline ctDNA scores tracked tumor burden but did not distinguish responders from non-responders, whereas ctDNA after one treatment cycle had a higher relative decrease in responders, six weeks before the next scan. We developed an ordinary differential equation model linking tumor volume to the ctDNA released by tumor cells. Sensitivity, identifiability, and model-selection analyses reduced it to two patient-specific parameters, proliferation and shedding rates, which reproduced individual volume and ctDNA trajectories, including a transient ctDNA spike at treatment onset. Proliferation was faster in non-responders. This framework supports model-based, ctDNA-guided monitoring of immunotherapy response.

## 1. Introduction

Anal squamous cell carcinoma (ASCC) is an aggressive malignancy characterized by limited systemic treatment options and dismal outcomes in the metastatic setting. Despite a low incidence rate of just two new cases per 100,000 people per year among the general US population, cases have been steadily rising by two to three percent per year, particularly among older adults and women^1^. Approximately 80–90% of ASCC cases are attributable to human papillomavirus (HPV) infection^2,3^. Standard of care treatment for non-metastatic disease includes concurrent chemoradiation^4^, while immunotherapy – including the anti-PD-1 antibody pembrolizumab – is used in the recurrent/metastatic setting^5,6^. HPV-associated cancers generally carry a better prognosis than HPV-negative cancers, attributed to enhanced immunogenicity ^7^ and increased treatment sensitivity^8^, which has motivated active investigation of de-escalation strategies in this disease^9^. However, response to single-agent immunotherapy in advanced ASCC remains modest and variable – 10% to 24% after failing initial chemotherapy^10^ and about 55.8% in combination^11^, and identifying which patients benefit from a given regimen remains uncertain^5,6^.

Like most solid tumors, ASCC treatment response is assessed through serial imaging; however, such surveillance is limited in that imaging occurs at infrequent intervals (every six to nine weeks) and is expensive and inconvenient. Moreover, the structural changes seen on imaging lag behind the underlying tumor biology, delaying assessment of a treatment’s efficacy. Consequently, a therapy’s effect may not be apparent until well after a meaningful biological change has occurred, and as such blood biomarkers like cell-free DNA (cfDNA) and circulating tumor DNA (ctDNA), can help capture the treatment effectiveness.

Most cfDNA in the bloodstream is derived from normal cells undergoing apoptosis, necrosis and active secretion, while a small proportion is derived from tumor cells and is known as circulating tumor DNA (ctDNA). ctDNA is released mainly by dying tumor cells but also through active secretion by viable cells^12^ **(Figure 1)**. Circulating tumor DNA (ctDNA) offers a complementary, minimally invasive window into tumor dynamics^13^. As ctDNA has a short circulating half-life that reflects near-real-time tumor activity ^14-16^ and can be collected frequently, ctDNA is an attractive candidate for real-time monitoring, detection of minimal residual disease (MRD), and early prediction of progression. HPV-associated cancers offer a compelling setting as tumor-specific viral sequences can be quantified with validated assays such as tumor tissue-modified viral HPV DNA (TTMV-HPV DNA), yielding an HPV-specific measure of tumor burden^2,17,18^. In a study of ASCC, TTMV-HPV DNA measured during routine pre- and post-treatment surveillance has shown high per-patient sensitivity and specificity for recurrence (82.8% and 98.4%, respectively) across 117 patients at seven U.S. centers. Positive tests were strongly associated with the presence of disease (positive predictive value, 96%) and demonstrated a median lead time of 59 days before clinical or radiographic confirmation ^18,19^.

Translating ctDNA measurements into clinically actionable predictions, however, requires more than observational correlation. The amount of ctDNA in a patient’s blood reflects several interacting processes: the number and proliferation of tumor cells, how much tumor DNA they release (largely through cell death, but also through active secretion), whether that DNA can reach the circulation through the tumor’s vasculature, and how quickly it is cleared^12,16,20^. Mathematical oncology – integration of mechanistic models with experimental and clinical data – provides a framework for capturing these dynamics and deriving patient-specific predictions^21,22^.Models of ctDNA dynamics have already shown clinical value. Avanzini et al. linked ctDNA levels to tumor size, establishing how large a lung tumor must be before it is detectable in blood^12^. In colorectal cancer, Khan et al. fit a two-population biexponential model to show how serial ctDNA modeling forecast treatment failure and progression before imaging did^23^. Janssen et al. also used nonlinear mixed-effects modeling of EGFR ctDNA to predict progression in lung cancer before imaging^24^, and Ribba et al. used a bi-exponential ctDNA model to show and inform the optimal ctDNA sampling times and proposed a joint tumor–ctDNA model with a shared decay parameter^25^. Longitudinal joint models have likewise shown that ctDNA trajectories predict progression-free or overall survival across regimens^26,27^, and Li et al. recently demonstrated *in silico* that early ctDNA kinetics differ by treatment mechanism and can serve as a dynamic biomarker of response, including a transient rise in ctDNA shortly after treatment initiation as therapy-induced cell death releases tumor DNA into circulation^28^. These models establish that ctDNA dynamics carry predictive information, but most are theoretical and simulation based. What remains less developed is a *coupled volume–ctDNA model that describes real, longitudinal, patient-specific data under treatment*.

The purpose of this study is to determine whether a simple mechanistic model can capture patient-specific longitudinal patterns of ctDNA and tumor volume in patients with ASCC receiving immunotherapy. We hypothesize that ctDNA *dynamics*, rather than a single value, can differentiate between those who will and will not continue to respond to treatment. We use ordinary differential equations (ODEs) to develop a coupled model, calibrate it to patient-specific data, and identify model-derived features that may serve as markers of treatment response. To our knowledge, this is the first mechanistic model coupling tumor volume and ctDNA to be calibrated to longitudinal, patient-specific clinical data in an HPV-associated cancer like ASCC.

## 2. Methods

### 2.1 Patient cohort and data

Longitudinal data were collected as part of a multicenter, single-arm phase II trial of pembrolizumab in metastatic or locally advanced incurable ASCC led by Dana-Farber Cancer Institute (NCT02919969)^6^. Patients received 200 mg of pembrolizumab intravenously every three weeks for up to 2 years. TTMV-HPV ctDNA (hereafter called ctDNA) was measured using the NavDx assay (CareDx, Inc., Brisbane, CA), a clinically validated assay for HPV-driven cancers^17^, at baseline and each cycle for the first three cycles, then every other cycle, while tumor burden was quantified as the sum of longest diameters (SLD) of target lesions, taken from the CT or MRI measurements recorded at each restaging scan in the trial and assessed by RECIST v1.1^29^ at baseline, treatment start, and approximately every nine weeks thereafter **(Figure 2)**.

**Figure 2.**
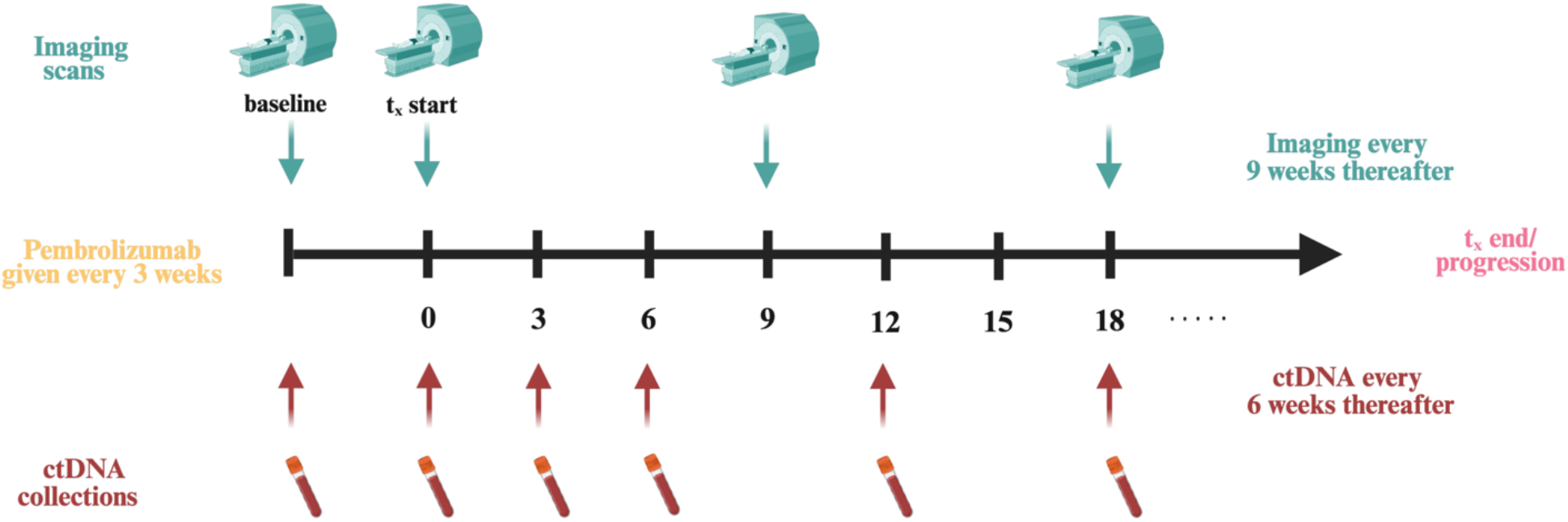
Serial ctDNA and imaging schedule in the pembrolizumab trial (NCT02919969) Patients received pembrolizumab 200 mg intravenously every 3 weeks. Blood for circulating tumor DNA (ctDNA) analysis was collected at every cycle for the first three cycles (weeks 0, 3, and 6) and every other cycle thereafter (weeks 12, 18, 24, …) until treatment discontinuation or disease progression, whichever occurred first (t_*x*_ end /progression). Imaging scans obtained at baseline and treatment start, every 9 weeks the discretion of the provider.

Of the 32 enrolled patients, 26 had both longitudinal RECIST and ctDNA data and were included in our analysis **(Figure S1)**. In the full trial population, the confirmed objective response rate was 9.4% and median progression-free survival was 2.2 months (median follow-up 12.6 months)^6^. For this analysis, patients were classified by their most recent RECIST 1.1 assessment^29^: those with complete response (CR, n = 1), partial response (PR, n = 4), or stable disease (SD, n = 9) were classified as responders (n = 14), and those with progressive disease were classified as non-responders (n = 12). Four of the patients with stable disease had unconfirmed SD. Because groups were assigned from the most recent assessment rather than from confirmed best overall response, response categories may differ from those reported for the trial **(Supplementary Data File 1**).

### 2.2 Baseline and early-dynamics analysis of ctDNA

Plasma TTMV-HPV DNA was measured with NavDx (CareDx, Inc., Brisbane, CA), a clinically validated assay for HPV-driven cancers^17^. Results are reported as a TTMV-HPV DNA score, where 0 indicates that no TTMV-HPV DNA was detected; throughout, ‘ctDNA’ refers to this score.

The relationship between ctDNA and tumor burden was characterized using two complementary analyses. First, we evaluated the relationship between baseline ctDNA and baseline tumor volume using Spearman’s rank correlation. Two patients had no baseline ctDNA sample and three had undetectable ctDNA at all timepoints, leaving 21 patients for this analysis. Second, to determine whether early changes in ctDNA differentiate patients by clinical outcome, we computed each patient’s log_10_ fold change, log_10_ 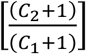 in ctDNA, over three intervals: baseline to Cycle 2, baseline to Cycle 3, and Cycle 2 to Cycle 3. To include undetectable results, 1 count was added to every TTMV-HPV DNA score before log transformation. Log fold changes were compared between responders and non-responders using the two-sided Mann-Whitney U (Wilcoxon rank-sum) test. Patients without a ctDNA sample at either timepoint of an interval were excluded from the corresponding analysis, as were patients whose ctDNA was undetectable at all timepoints. The number of patients in each comparison is given in **Supplementary Data File 1**.

### 2.3 Mathematical model

We developed a mathematical model of tumor volume (*V*), quantitative ctDNA score (*C*), and treatment (γ) dynamics **(Figure 3)**. In the absence of treatment, tumor volume is governed by the natural balance between tumor cell birth (*b*) and death (*d*), with ctDNA shed from dying cells at rate *q* and cleared from circulation at rate ε.

**Figure 3.**
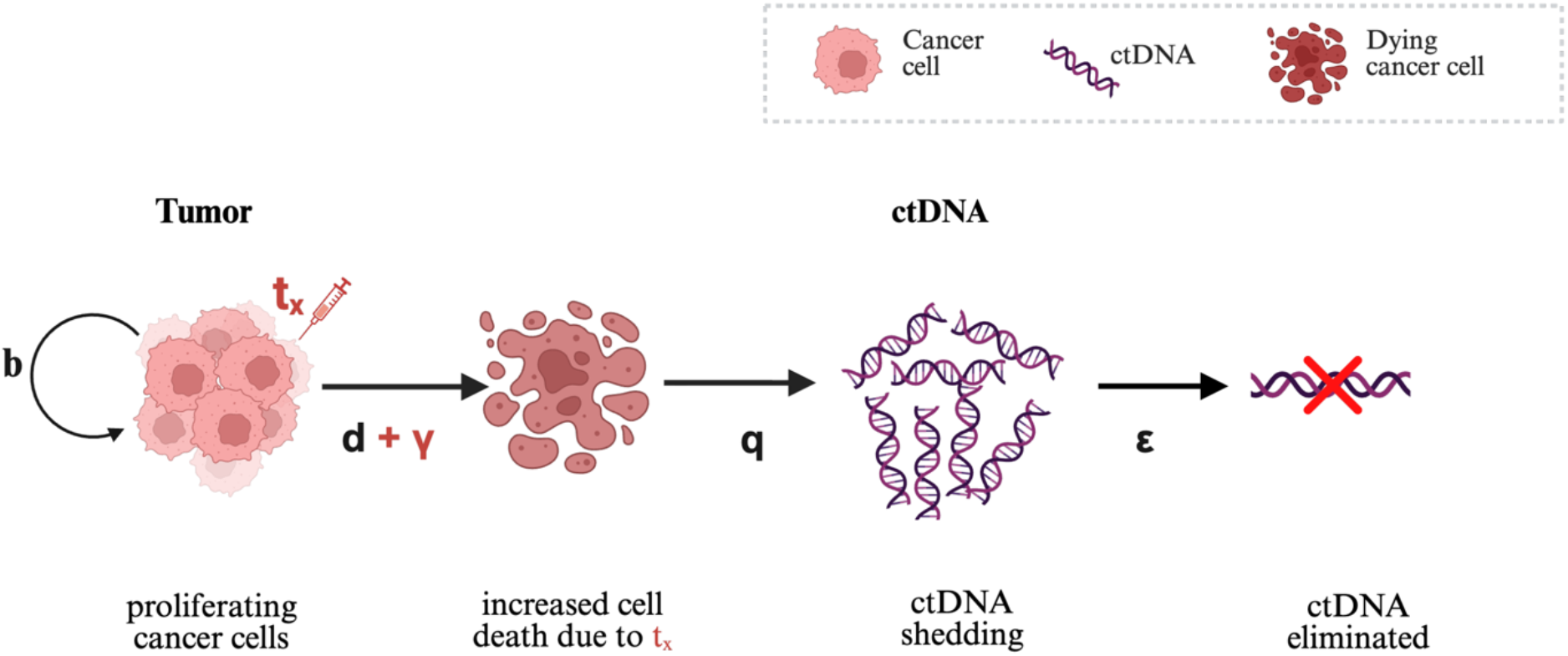
Model Schematic. Tumor cells proliferate (*b*) and are lost through natural death (*d*), plus a treatment-induced term (γ) that is active while the patient is on treatment (*t*_*x*_). Dying cells shed circulating tumor DNA (ctDNA) into the bloodstream at rate *q*, and ctDNA is cleared from circulation at rate ε. Treatment sensitivity (γ) decays over time at a rate δ, allowing the model to capture a waning response.

Mathematically, this is described using ODEs as:

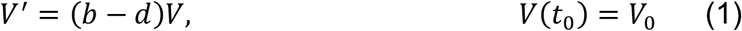

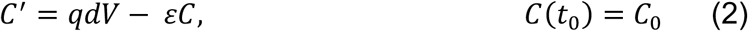

Motivated by the Claret tumor growth inhibition (TGI) model^30,31^, in the presence of pembrolizumab treatment *t*_*x*_, volume is reduced at rate γ, while the treatment effect γ decays at rate δ as resistance develops. This contributes to additional dead tumor cells producing ctDNA. Mathematically, the model is given as

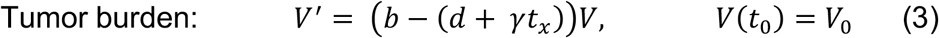

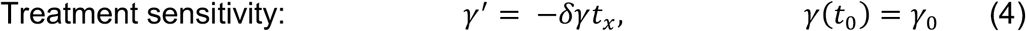

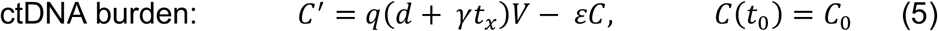

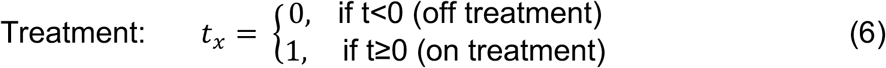

where, *t*_*x*_ denotes the Heaviside step function indicating whether or not the protocol therapy has been initiated. The parameters, including initial conditions, are described in **(Table 1)**. Because treatment adds a death term (γ*t*_*x*_) that switches on at treatment onset, the model predicts an abrupt increase in tumor-cell death and therefore in ctDNA shedding. This produces a transient post-treatment ctDNA spike that resolves as tumor volume falls, consistent with the early surge described by Li et al.^28^.

**Table 1.**
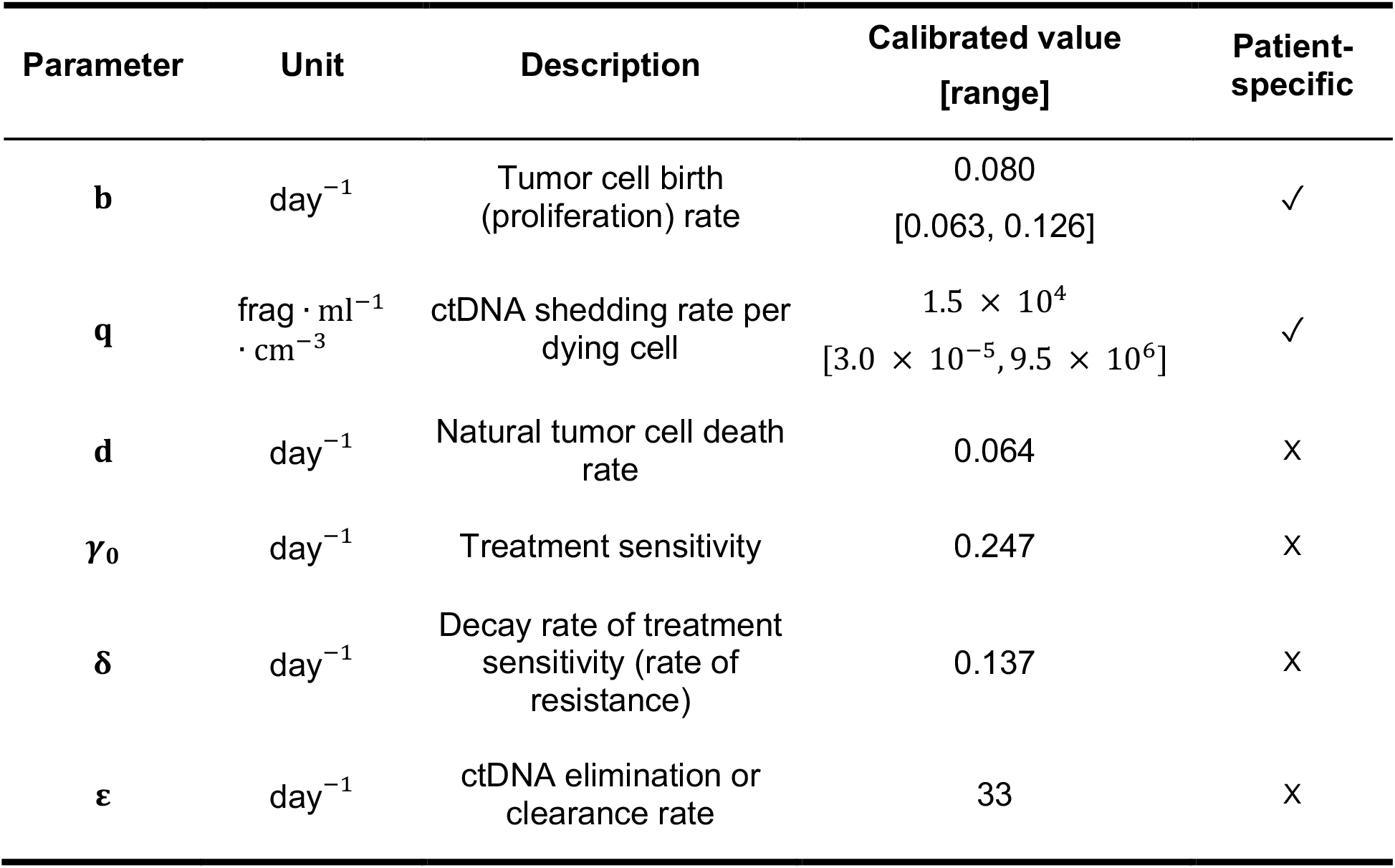
Calibrated model parameters. Parameter estimates for the coupled ODE model of tumor volume and ctDNA dynamics, obtained by fitting data from the NCT02919969 cohort. All rate parameters are reported in day^−1^; the shedding parameter *q*, which converts tumor volume to ctDNA concentration, carries units of frag · ml^−1^ · cm^−3^. For parameters estimated on a per-patient basis, the calibrated value shown is the cohort median with the minimum and maximum across patients shown in brackets. The remaining parameters were fixed at population-level values. The final column indicates whether each parameter was fitted individually per patient (✓) or held constant across the cohort (X).

| Parameter | Unit | Description | Calibrated value<br>[range] | Patient-specific |
|---|---|---|---|---|
| **b** | day<sup>−1</sup> | Tumor cell birth (proliferation) rate | 0.080<br>[0.063, 0.126] | ✓ |
| **q** | frag · ml<sup>−1</sup> · cm<sup>−3</sup> | ctDNA shedding rate per dying cell | 1.5 × 10<sup>4</sup><br>[3.0 × 10<sup>−5</sup>, 9.5 × 10<sup>6</sup>] | ✓ |
| **d** | day<sup>−1</sup> | Natural tumor cell death rate | 0.064 | X |
| **γ<sub>0</sub>** | day<sup>−1</sup> | Treatment sensitivity | 0.247 | X |
| **δ** | day<sup>−1</sup> | Decay rate of treatment sensitivity (rate of resistance) | 0.137 | X |
| **ε** | day<sup>−1</sup> | ctDNA elimination or clearance rate | 33 | X |

### 2.4 Sensitivity and Identifiability Analysis

Prior to calibrating the model to the data, local sensitivity analysis ^32 33^ was performed to assess the variation in the model output in response to small perturbations in the model parameters. The sensitivity matrix is given by

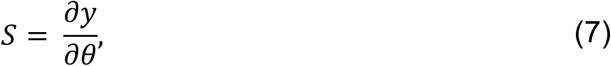

where *y* = [*V, C*] is the model output and *θ* = {*γ*_0_, *b, d, q, ϵ*} are the model parameters. The sensitivities were computed numerically ^32^ and then ranked by averaging over time using the two-norm, ordering parameters from most to least sensitive. Next, correlation analysis was used to determine parameter identifiability, that is whether or not a parameter could be uniquely determined ^34^. Sensitive parameters were analyzed for pairwise correlations using the correlation method ^32,35^. Given the covariance matrix *C* = *A*^−1^ = (*S*^*T*^*S*)^−1^, the correlation coefficients are given by

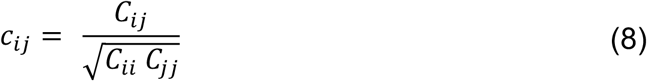

A parameter pair (*i, j*) is considered correlated when | *c*_*ij*,_ | > ξ→1. Such pairs can compensate for one another without changing the model fit and therefore cannot be estimated independently. We chose a strict threshold, ξ = 0.99, to flag only near-redundant parameter pair.

### 2.5 Parameter estimation and model selection

We estimated patient-specific parameters by minimizing the least-squares error between the model output and data. We assessed the fits to both outputs (*V* and *C*) using the root-mean-square error (RMSE) and the normalized RMSE (nRMSE). As our primary goal was to determine specific markers of treatment response, we sought to reduce the number of patient-specific parameters while maintaining adequate model fit. Starting from a fully patient-specific fit, we iteratively fixed individual parameters to a shared cohort value (typically the mean estimated from the preceding fit) and refit the remaining patient-specific parameters. Parameter distributions, coefficients of variation (*CV* = standard deviation/mean), and model fit were evaluated at each step to determine which parameters could be shared across patients and which should remain patient-specific. Candidate model structures were then compared using the Akaike Information Criterion (AIC)^36,37^, given by

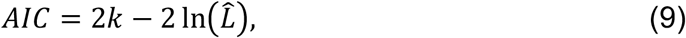

where *k* is the number of estimated parameters and 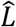 is the maximized likelihood. For each candidate structure we also assessed whether the responder vs. non-responder differences in the optimized parameters were statistically significant. Responder vs. non-responder differences were tested with the Mann-Whitney U (Wilcoxon rank-sum) test (α = 0.05).

## 3. Results

### 3.1 Baseline ctDNA correlates with tumor burden but does not stratify response

Baseline ctDNA score ranged from undetectable to about 10^6^, roughly six orders of magnitude among detectable samples, while baseline SLD ranged from approximately 14 to 152mm. Spearman rank correlation showed a statistically significant positive association between the two (ρ = 0.67, *p* = .001, *n* = 21) **(Figure 4a)**. Thus, patients with larger tumors at baseline had higher baseline ctDNA levels. However, baseline measurements did not separate responders from non-responders. Although responders tended to have somewhat lower baseline SLD, the two distributions overlapped substantially between responders and non-responders, and baseline ctDNA levels did not differ significantly between the two groups (*p* = .093). Thus, while baseline ctDNA correlates with baseline tumor burden, a single pre-treatment measurement is insufficient on its own to determine whether a patient will respond to immunotherapy treatment.

**Figure 4.**
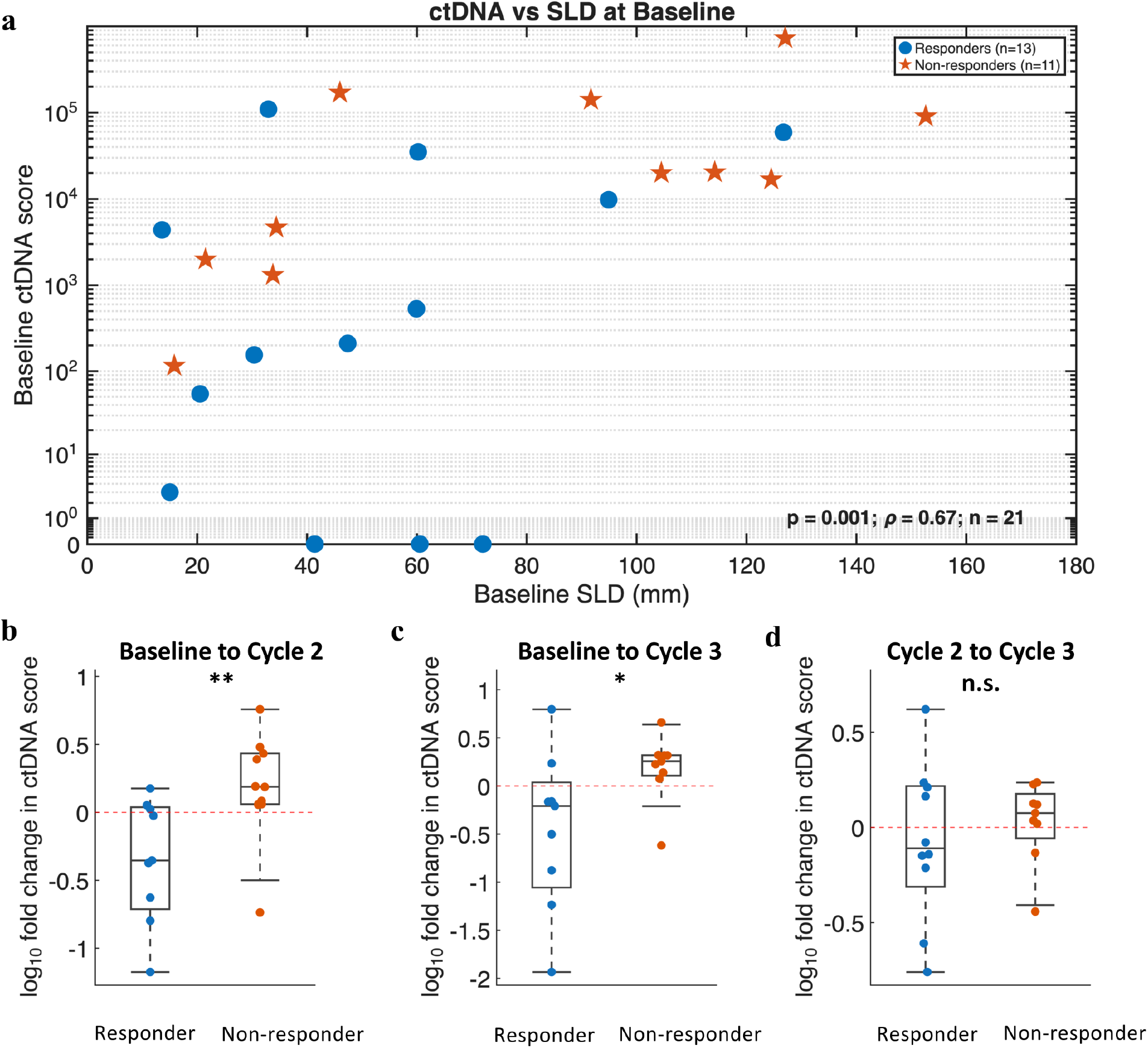
Circulating tumor DNA dynamics and correlation with tumor burden in responders versus non-responders. (A) Scatter plot showing baseline ctDNA versus sum of longest tumor diameters (SLD). ctDNA correlated with tumor size (Spearman ρ = 0.67, *p* < 0.001). Statistical comparisons by Mann-Whitney U test. (B-D) Box plots comparing log_10_ fold change in ctDNA between responders (blue) and non-responders (orange) from (B) baseline to Cycle 2 (C2D1), (C) baseline to Cycle 3 (C3D1), and (D) Cycle 2 to Cycle 3. Red crosses mark group means. Changes from baseline to Cycle 2 (** *p* < 0.01) and from baseline to Cycle 3 (* *p* < 0.05) differed significantly between groups; the change from Cycle 2 to Cycle 3 did not.

### 3.2 Early change in ctDNA score distinguishes responders from non-responders

We next investigated if ctDNA dynamics, rather than static measurements from baseline could stratify patients by treatment response. Between baseline and Cycle 2, after a single cycle of pembrolizumab, responders saw a larger decline in ctDNA compared to non-responders. The median log_10_ fold change was −0.35 vs. 0.19, corresponding to a roughly 0.44-fold decrease vs. a 1.5-fold increase (*n* = 9 and 11; *p* = .004; **Figure 4b**). Ten of 11 non-responders had rising ctDNA over this interval. Only 3 of 9 responders did, and none of those increases exceeded 1.5-fold. This pattern is consistent with treatment-induced ctDNA clearance in responders and continued shedding from growing tumors in those who do not. It was evident at Cycle 2, approximately six weeks before the first scheduled CT assessment (around Week 9).

Between baseline and Cycle 3, the difference persisted (median −0.21 vs. 0.26; *n* = 9 and 9; *p* = .042; **Figure 4c**): the ctDNA score rose in 8 of 9 non-responders compared with 2 of 9 responders. Between Cycle 2 and Cycle 3, ctDNA changed little in either group (median −0.11 vs. 0.08; *n* = 10 and 9; *p* = .49, *n. s*.; **Figure 4d**), indicating that the separation arises mainly from the initial change between baseline and Cycle 2.

Together with the baseline analysis, these results indicate that the early trajectory of ctDNA carries outcome information and its pre-treatment level does not. Because this signal is a change over time rather than a static value, it motivates the coupled volume– ctDNA model developed below.

### 3.3 Coupled volume-ctDNA model accurately describes patient-specific dynamics

Local sensitivity analysis ranked the tumor proliferation rate *b* as the most sensitive parameter and the rate of resistance δ as the least sensitive. Correlation analysis found correlations between *b* and *d*, which affect tumor volume only through their net difference (*b* − *d*) and with ε and *q*, which affect ctDNA shedding and clearance. After accounting for these, the sensitive and identifiable parameters were [*γ*_0_, *b, δ, q*].

Given this, the model was calibrated to the patient-specific longitudinal ctDNA and volume data, where [*γ*_0_, *b, δ, q*] were allowed to be patient-specific and *d* and ϵ were fixed (*d* = 0.064, ϵ = 33). After evaluating the model fits using the nRMSE, coefficients of variation for each parameter **(Figure S5)**, and the AIC value, we sequentially reduced the model complexity by allowing a subset of the parameters to be patient-specific and the remaining parameters to be uniform **(Supplementary Data File 1-S3, S4, S5, Supplementary Figures S3, S4, S5)**. This was done to not only reduce complexity, but also to avoid model overfitting. Comparisons between the quality of the fits to both ctDNA and volume, as well as the AIC value **(Table S1)**, found that allowing the tumor proliferation rate *b* and the ctDNA shedding rate *q* to be patient-specific, while the remaining parameters *γ*_0_, *δ, d*, and ϵ to be uniform between patients, resulted in the best fits to the data.

The model was able to describe both ctDNA and volume dynamics in responders and non-responders **(Figure 5a)**. Patients 1 and 7 in **Figure 5** are shown as representative examples of the dominant ctDNA pattern in each group: a fall in ctDNA after the first cycle in responders, and a continued rise in non-responders. In Patient 1 (responder), volume fell sharply after treatment onset and then regrew slowly, remaining below baseline through Day 180. ctDNA showed an initial post-treatment spike, followed by a decline and gradual regrowth. In Patient 7 (non-responder), though both ctDNA and volume initially decline, regrowth resumed in both within about three weeks, and both volume and ctDNA exceeded baseline by Day 60.

**Figure 5.**
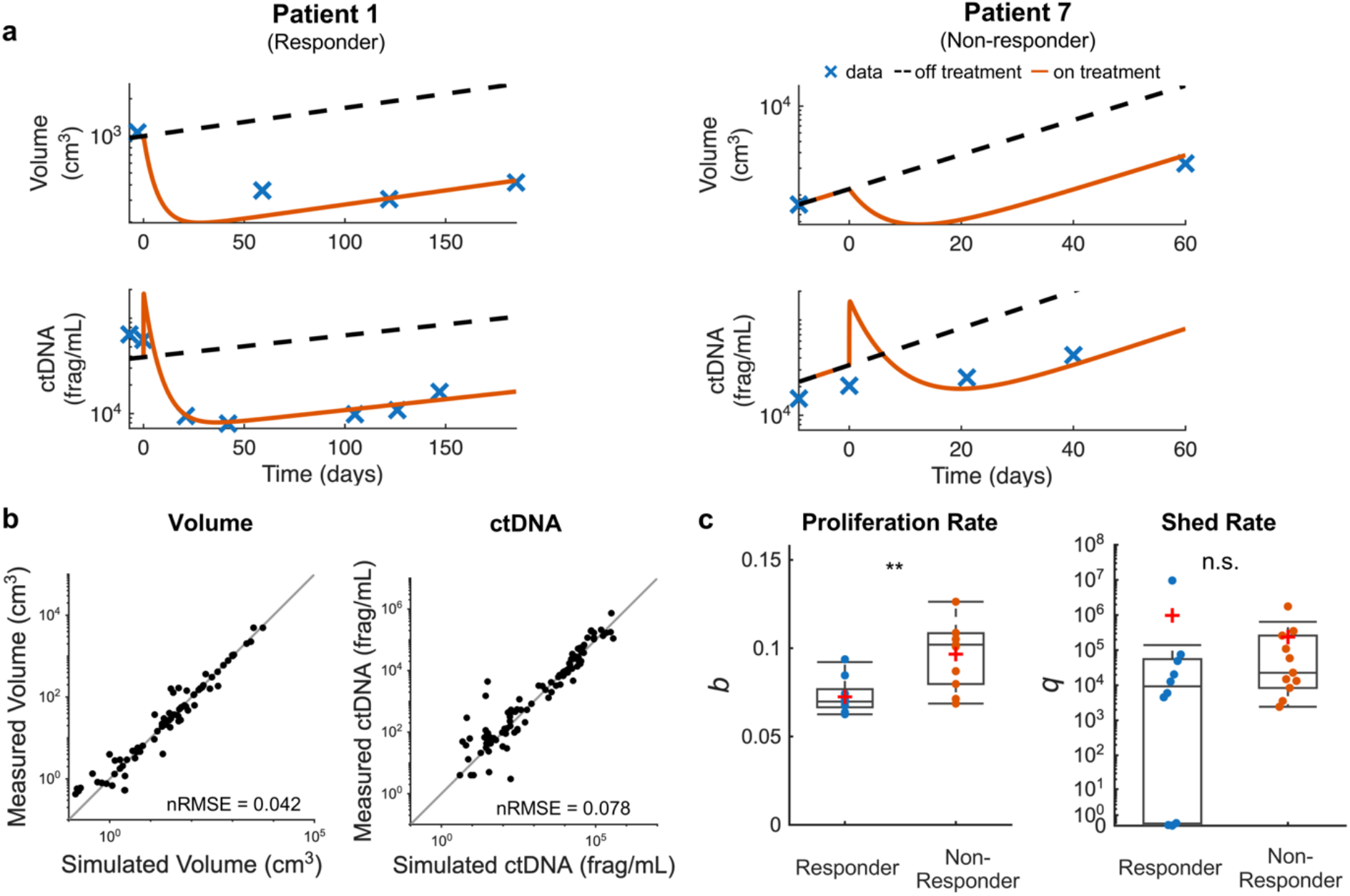
Patient-specific model calibration and parameter differences between responders and non-responders. (A) Representative fits from the selected model (Model C; see Table 2) for a responder (Patient 1) and non-responder (Patient 7), showing tumor volume (cm^3^) and ctDNA (frag/mL) on log axes over time. Crosses are measured data; red curves are on-treatment model fits; dashed curves are the model-simulated off-treatment trajectories. (B) Measured versus simulated volume (nRMSE = 0.042) and ctDNA (nRMSE = 0.078); dashed line is the line of identity. (C) The two patient-specific parameters compared between responders (blue) and non-responders (orange): growth rate *b* differed significantly (**, p < 0.01), shed rate *q* did not (n.s.). Mann-Whitney U test.

Comparisons between the simulated and measured values for ctDNA and volume over all patients demonstrated that the model accurately described dynamics for the population as a whole (*nRMSE*_*V*_ = 0.042, *nRMSE*_*C*_ = 0.078) **(Figure 5b)**. The model was calibrated to the 21 patients with a usable longitudinal ctDNA series (10 responders, 11 non-responders). Parameter comparisons between responders and non-responders showed that non-responders had a significantly higher proliferation rate *b* than responders (median *b* = 0.06978 for responders, *b* = 0.102 for non-responders, *p* = 0.0028). Similar analysis for the ctDNA shed rate *q* showed that the values did not differ significantly between response groups. It is important to note that *b* was significantly higher in non-responders across all optimization structures **(Supplementary_Data_File 1 - File 6)**. As such, we can conclude that the proliferation rate is a significant driver of response in these patients.

## 4. Discussion

In this study, we asked whether a simple mechanistic model could describe the coupled dynamics of tumor volume and circulating TTMV-HPV DNA in patients with HPV-associated ASCC treated with pembrolizumab. Before modeling, we examined what the ctDNA measurements alone could tell us. Baseline ctDNA scores correlated with tumor burden (ρ = 0.67) but did not distinguish responders and non-responders. The early change did: between baseline and Cycle 2, after a single cycle, ctDNA fell in 6 of 9 responders and rose in 10 of 11 non-responders (*p* = .004). This separation appeared about six weeks before the first scheduled restaging scan.

The baseline correlation we observed is consistent with reports in other HPV-associated cancers, but published estimates vary. In advanced anal carcinoma, Bernard-Tessier et al. reported a weaker association between HPV ctDNA and tumor burden (ρ = 0.32)^5^, and in a pan-cancer cohort treated with pembrolizumab, baseline ctDNA did not correlate significantly with RECIST target-lesion size^38^. These differences likely reflect the factors that determine how much tumor DNA reaches the blood, including cell turnover, tumor vascularity, and site of disease. They may also reflect when baseline was measured: here, before salvage immunotherapy, whereas other studies sampled at other points in the disease course, such as before first-line chemotherapy^5^. A single ctDNA value is therefore an imperfect proxy for tumor size, which is part of why we focused on its dynamics. Hence the motivation for a dynamic description: the ctDNA response signal lives in the early trajectory.

Our early-change finding parallels that of Bratman et al., who measured personalized ctDNA in 94 patients with five types of advanced solid tumors treated with pembrolizumab in the INSPIRE trial^38^. A decrease in ctDNA from baseline to the start of Cycle 3 was associated with objective response and with longer overall and progression-free survival, and all 12 patients who cleared ctDNA were alive at last follow-up. Our study differs in three ways. It is restricted to HPV-associated ASCC and uses a viral DNA assay that does not require tumor sequencing rather than a tumor-informed mutation panel. The ctDNA decline separated groups earlier, at Cycle 2 after a single cycle. And our outcome was response, as defined here rather than survival, so whether the Cycle 2 change also predicts survival in ASCC remains to be tested. These observations motivated a mechanistic description: the response signal sits in the early trajectory, which a single measurement cannot capture.

The mathematical model reproduced both tumor volume and ctDNA across the cohort, and the model-reduction was informative beyond fit quality. Given our sensitivity and identifiability analysis, and coefficient of variations **(Figures S3, S4, and S5)**, we initially fixed *d* and ε and as kept the remaining parameters patient-specific. Even with the treatment-sensitivity parameters *γ*_0_ and *δ* shared across the cohort, a structure retaining only the proliferation rate *b* and the shedding rate *q* per patient, Structure 3 still reproduced both outputs (*nRMSE*_*V*_ = 0.042, *nRMSE*_*C*_ = 0.078). The fully patient-specific model fit best and had the lowest AIC, as expected of the least-constrained structure; we nonetheless adopted the most reduced model as our working structure because it describes the whole cohort with only two patient-specific parameters and localizes the between-patient differences to proliferation (*b*) and shedding (*q*), with all treatment-response parameters shared. This concentration of the predictive signal in patient-specific *dynamic* parameters, rather than baseline values, echoes findings from adaptive-therapy modeling in other cancers ^39,40^.

Across every candidate model structure, the proliferation rate b was the one parameter that consistently differed between responders and non-responders **(Table S1)**. This held even when *γ*_0_ and *δ* were free to vary by patient, so the difference is not an artifact of sharing the treatment-sensitivity parameters. Because volume depends only on the net rate (*b* − *d*) and d was fixed, this should be read as a difference in *net* tumor growth. The model therefore attributes differences in outcome mainly to how quickly tumors grow, rather than to how strongly they respond to treatment or how much ctDNA they shed. Both net growth and shedding reflect properties of individual tumors, so variation across patients is biologically expected.

Beyond model accuracy, the framework supports the clinical use for early, blood-based read-outs of response and carries real stakes for use in ASCC. HPV-associated cancers generally respond well to radiation and chemotherapy, which has motivated active investigation of de-escalation strategies to prevent overtreatment ^9^, while in the recurrent/metastatic setting of ASCC treated with immunotherapy, response rates remain modest and resistance can emerge after an initial benefit, making early identification of non-responders equally important ^5,6^. Consistent with this, real-world surveillance data show that TTMV-HPV DNA frequently anticipates clinical or radiographic recurrence ^19^. Whereas prior ctDNA models have been largely theoretical or simulation based ^12,23-25,28^, our model is, to our knowledge, the first coupled volume-ctDNA model calibrated to longitudinal patient data in HPV-associated ASCC, and it provides a mechanistic account of how these dynamics arise and a starting point for the early, model-based response monitoring that these data motivate.

We note a few limitations. First, the cohort is small (*n* = 26 from a single trial NCT02919969), and the responder/non-responder comparisons are correspondingly underpowered; a non-significant difference – such as that in the shedding rate *q* – should be read as inconclusive rather than as evidence of no difference, and by same token, the between-group differences that did reach significance (e.g., in the birth rate *b*) require confirmation in a larger cohort before they can be considered robust. Secondly, tumor volume is sampled sparsely – at baseline, treatment start, and roughly every nine weeks thereafter – so most patients contribute only a handful of volume measurements relative to the number of estimated parameters. Blood sampling also became sparser after Cycle 2 – 18 patients contributed to the baseline-to-Cycle 3 comparison versus 20 at Cycle 2, so the Cycle 3 comparison rests on fewer patients.. The model thus has substantial freedom to interpolate, and the low aggregate error reflects descriptive goodness-of-fit rather than predictive validation. Thirdly, because *b* and *d* are non-identifiable from volume, the localization of the response difference to proliferation should be understood as a statement about net growth, not a resolution of birth versus death. Finally, the model idealizes treatment as a binary on/off term without explicit pharmacokinetics or dose dependence and represents resistance as a monotonic decay of sensitivity that cannot recover, precluding re-sensitization or intermittent response.

Building on this descriptive foundation, few directions would move the framework from description toward prediction. Validation in an independent, more frequently sampled cohort would test whether the fits reflect genuine dynamics rather than interpolation of sparse data; formal uncertainty quantification would separate well-constrained patient-specific estimates from those limited by their bounds; and testing whether early-cycle model quantities anticipate time to progression (TTP) in held-out patients would be the natural next step, one not attempted here. Natural model extensions include replacing the binary treatment term with explicit drug pharmacokinetics, allowing treatment sensitivity to recover, and adapting the framework to the chemoradiotherapy setting in non-metastatic disease. More broadly, a validated model of this kind could allow clinicians to read out an early, blood-based estimate of whether a therapy is working, supporting model-guided treatment adaptation well before imaging would reveal progression ^21^.

## 5. Conclusion

In a cohort of patients with HPV-associated ASCC we were able to demonstreat the utility of ctDNA monitoring during treatment with pembrolizumab. Baseline ctDNA correlated with tumor burden but did not distinguish responders from non-responders. In contrast, the early change in ctDNA did: between baseline and Cycle 2 did: ctDNA declined in responders and rose in non-responders. Moreover, these changes were seen approximately six weeks before the first scheduled imaging. Next, our selected mechanistic model coupling tumor volume, treatment sensitivity, and ctDNA captured patient-specific dynamics across the cohort using only two patient-specific parameters, the proliferation rate *b* and ctDNA shedding rate *q*. Lastly, of these, the tumor proliferation rate b differed significantly between responders and non-responders, which places the difference in outcome in net tumor growth rather than in treatment sensitivity or ctDNA shedding. Together, these results show that the information about response lies in how ctDNA changes over time, not in any single measurement, and they support further development of model-based, blood-based response monitoring in ASCC over time.

## 6. Acknowledgments

The work of James Cleary is supported by the Donna V. Toelke Fund for Cancer Research.

## 7. Author Contributions

Model conception and development: Phebe M. Havor, Renee Brady-Nicholls Financial support: Renee Brady-Nicholls

Collection and assembly of clinical data: Brandon Huffman, James Cleary, James Jabalee

Data analysis and interpretation: All authors Manuscript writing: All authors

Final approval of manuscript: All authors Accountable for all aspects of the work: All authors

## 9. Data Availability

Data availability upon request.

## References

1 Siegel, R. L., Kratzer, T. B., Giaquinto, A. N., Sung, H. & Jemal, A. Cancer statistics, 2025. CA Cancer J Clin 75, 10–45 (2025). 10.3322/caac.21871

2 Andrioaie, I. M. et al. The Clinical Utility of Circulating HPV DNA Biomarker in Oropharyngeal, Cervical, Anal, and Skin HPV-Related Cancers: A Review. Pathogens 12 (2023). 10.3390/pathogens12070908

3 Morris, V. K. Circulating Tumor DNA in Advanced Anal Cancer: A Blood Biomarker Goes Viral. Clin Cancer Res 25, 2030–2032 (2019). 10.1158/1078-0432.Ccr-18-3982

4 Araradian, C., Walsh, M., Standage, H. & Tsikitis, V. L. Advances in the Management, Treatment, and Surveillance of Anal Squamous Cell Cancer. Cancers (Basel) 17 (2025). 10.3390/cancers17081289

5 Bernard-Tessier, A. et al. Clinical Validity of HPV Circulating Tumor DNA in Advanced Anal Carcinoma: An Ancillary Study to the Epitopes-HPV02 Trial. Clin Cancer Res 25, 2109–2115 (2019). 10.1158/1078-0432.Ccr-18-2984

6 Huffman, B. M. et al. Biomarkers of pembrolizumab efficacy in advanced anal squamous cell carcinoma: analysis of a phase II clinical trial and a cohort of long-term responders. J Immunother Cancer 12 (2024). 10.1136/jitc-2023-008436

7 Lechner, M., Liu, J., Masterson, L. & Fenton, T. R. HPV-associated oropharyngeal cancer: epidemiology, molecular biology and clinical management. Nature Reviews Clinical Oncology 19, 306–327 (2022). 10.1038/s41571-022-00603-7

8 Petrelli, F., Ghilardi, M., De Stefani, A., Nardone, M. & Capriotti, V. State of the Art of Systemic Therapy in HPV-Positive Oropharyngeal Squamous Cell Carcinoma: A Scoping Review. Diseases 14 (2026). 10.3390/diseases14020046

9 Wu, C., Kuzmin, P. & Julian, R. De-Escalation Strategies in HPV-Associated Oropharynx Cancer: A Historical Perspective with Future Direction. Cancers (Basel) 16 (2024). 10.3390/cancers16152733

10 Jayakrishnan, T., Yadav, D., Huffman, B. M. & Cleary, J. M. Immunological Checkpoint Blockade in Anal Squamous Cell Carcinoma: Dramatic Responses Tempered By Frequent Resistance. Curr Oncol Rep 26, 967–976 (2024). 10.1007/s11912-024-01564-z

11 Dhawan, N., Afzal, M. Z. & Amin, M. Immunotherapy in Anal Cancer. Curr Oncol 30, 4538–4550 (2023). 10.3390/curroncol30050343

12 Avanzini, S. et al. A mathematical model of ctDNA shedding predicts tumor detection size. Sci Adv 6 (2020). 10.1126/sciadv.abc4308

13 Huang, X., Deng, Y. & Shen, Y. Circulating tumor DNA in neoadjuvant therapy for solid tumors. Frontiers in Oncology Volume 16 – 2026 (2026). 10.3389/fonc.2026.1909934

14 Bartolomucci, A. et al. Circulating tumor DNA to monitor treatment response in solid tumors and advance precision oncology. NPJ Precis Oncol 9, 84 (2025). 10.1038/s41698-025-00876-y

15 Pessoa, L. S., Heringer, M. & Ferrer, V. P. ctDNA as a cancer biomarker: A broad overview. Crit Rev Oncol Hematol 155, 103109 (2020). 10.1016/j.critrevonc.2020.103109

16 Sanz-Garcia, E., Zhao, E., Bratman, S. V. & Siu, L. L. Monitoring and adapting cancer treatment using circulating tumor DNA kinetics: Current research, opportunities, and challenges. Sci Adv 8, eabi8618 (2022). 10.1126/sciadv.abi8618

17 Gunning, A. et al. Analytical Validation of NavDx, a cfDNA-Based Fragmentomic Profiling Assay for HPV-Driven Cancers. Diagnostics (Basel) 13 (2023). 10.3390/diagnostics13040725

18 Kabarriti, R. et al. Evaluating Tumor Tissue Modified Viral (TTMV)-HPV DNA for the Early Detection of Anal Squamous Cell Carcinoma Recurrence. Cancers (Basel) 17 (2025). 10.3390/cancers17020174

19 Kabarriti, R. et al. Resolving Clinically Indeterminate Findings During Anal Cancer Surveillance with TTMV-HPV DNA. Cancers (Basel) 18 (2025). 10.3390/cancers18010035

20 Zheng, G. When biology limits the biomarker: factors influencing ctDNA shedding. ASCO Daily News (2026).

21 Enderling, H., Alfonso, J. C. L., Moros, E., Caudell, J. J. & Harrison, L. B. Integrating Mathematical Modeling into the Roadmap for Personalized Adaptive Radiation Therapy. Trends in cancer 5, 467–474 (2019). 10.1016/j.trecan.2019.06.006

22 Scibilia, K. R. et al. Mathematical Oncology: How Modeling Is Transforming Clinical Decision-Making. Cancer Res 85, 4866–4879 (2025). 10.1158/0008-5472.Can-25-0750

23 Khan, K. H. et al. Longitudinal Liquid Biopsy and Mathematical Modeling of Clonal Evolution Forecast Time to Treatment Failure in the PROSPECT-C Phase II Colorectal Cancer Clinical Trial. Cancer Discov 8, 1270–1285 (2018). 10.1158/2159-8290.Cd-17-0891

24 Janssen, J. M. et al. Longitudinal nonlinear mixed effects modeling of EGFR mutations in ctDNA as predictor of disease progression in treatment of EGFR-mutant non-small cell lung cancer. Clin Transl Sci 15, 1916–1925 (2022). 10.1111/cts.13300

25 Ribba, B., Roller, A., Helms, H. J., Stern, M. & Bleul, C. Circulating tumor DNA: Opportunities and challenges for pharmacometric approaches. Front Pharmacol 13, 1058220 (2022). 10.3389/fphar.2022.1058220

26 Johnson, M. et al. Longitudinal Circulating Tumor DNA Modeling to Predict Disease Progression in First-Line Mutant Epidermal Growth Factor Receptor Non-Small Cell Lung Cancer. Clin Pharmacol Ther 115, 349–360 (2024). 10.1002/cpt.3113

27 Yuan, M. et al. Treatment-agnostic joint modeling of longitudinal circulating tumor dna predicts survival across first-line regimens in metastatic non-squamous NSCLC. Eur J Pharm Sci 214, 107275 (2025). 10.1016/j.ejps.2025.107275

28 Li, A., Lou, E., Leder, K. & Foo, J. Early Circulating Tumor DNA Kinetics as a Dynamic Biomarker of Cancer Treatment Response. JCO Clin Cancer Inform 9, e2400160 (2025). 10.1200/cci-24-00160

29 Eisenhauer, E. A. et al. New response evaluation criteria in solid tumours: revised RECIST guideline (version 1.1). Eur J Cancer 45, 228–247 (2009). 10.1016/j.ejca.2008.10.026

30 Claret, L. et al. Model-based prediction of phase III overall survival in colorectal cancer on the basis of phase II tumor dynamics. J Clin Oncol 27, 4103–4108 (2009). 10.1200/jco.2008.21.0807

31 Glazar, D. J. et al. Early response dynamics predict treatment failure in patients with recurrent and/or metastatic head and neck squamous cell carcinoma treated with cetuximab and nivolumab. Oral Oncol 127, 105787 (2022). 10.1016/j.oraloncology.2022.105787

32 Olufsen, M. S. & Ottesen, J. T. A practical approach to parameter estimation applied to model predicting heart rate regulation. Journal of mathematical biology 67, 39–68 (2013). 10.1007/s00285-012-0535-8

33 Pope, S. R. et al. Estimation and identification of parameters in a lumped cerebrovascular model. Math Biosci Eng 6, 93–115 (2009). 10.3934/mbe.2009.6.93

34 Miao, H., Xia, X., Perelson, A. S. & Wu, H. ON IDENTIFIABILITY OF NONLINEAR ODE MODELS AND APPLICATIONS IN VIRAL DYNAMICS. SIAM Rev Soc Ind Appl Math 53, 3–39 (2011). 10.1137/090757009

35 Ottesen, J. T., Mehlsen, J. & Olufsen, M. S. Structural correlation method for model reduction and practical estimation of patient specific parameters illustrated on heart rate regulation. Math Biosci 257, 50–59 (2014). 10.1016/j.mbs.2014.07.003

36 Akaike, H. A new look at the statistical model identification. IEEE Transactions on Automatic Control 19, 716–723 (1974). 10.1109/TAC.1974.1100705

37 Burnham, K. P. a. A. D.R. Model Selection and Inference: A Practical Information-Theoretic Approach. 2nd Edition edn, (2002).

38 Bratman, S. V. et al. Personalized circulating tumor DNA analysis as a predictive biomarker in solid tumor patients treated with pembrolizumab. Nat Cancer 1, 873–881 (2020). 10.1038/s43018-020-0096-5

39 Brady-Nicholls, R. et al. Predicting patient-specific response to adaptive therapy in metastatic castration-resistant prostate cancer using prostate-specific antigen dynamics. Neoplasia 23, 851–858 (2021). 10.1016/j.neo.2021.06.013

40 Zhang, J., Cunningham, J. J., Brown, J. S. & Gatenby, R. A. Integrating evolutionary dynamics into treatment of metastatic castrate-resistant prostate cancer. Nat Commun 8, 1816 (2017). 10.1038/s41467-017-01968-5

