## Supplemental Document for "Coupled Modeling of Tumor Volume and ctDNA Dynamics in HPV-Associated Anal Squamous Cell Carcinoma"

**Affiliations:**
[1]Integrated Mathematical Oncology Department, H. Lee Moffitt Cancer Center & Research Institute, Tampa, Florida
[2]Navaris, Inc., Waltham, Massachusetts
[3]Department of Medical Oncology, Dana- Farber Cancer Institute, Boston, Massachusetts
[4]Department of Medicine, Harvard Medical School, Boston, Massachusetts

The PDF file includes:

Notes S1 to S4

Figs. S1 to S5

Table S1

Data File S1 (separate Excel file)

References

**Note S1. Patient cohort and response classification**

Longitudinal tumor and ctDNA data were obtained from a multicenter, single-arm phase II trial of pembrolizumab in advanced anal squamous cell carcinoma (NCT02919969)[1]. Of 32 enrolled patients, five had no evaluable RECIST assessment (early discontinuation or missing scans) and one had no ctDNA measurement, leaving 26 patients with both longitudinal RECIST and ctDNA data (Figure S1).

Patients were classified by their most recent RECIST 1.1 assessment[3]. Patients with complete response (CR, n = 1), partial response (PR, n = 4), or stable disease (SD, n = 9) were classified as responders (n = 14), and patients with progressive disease (PD) were classified as non-responders (n = 12). Four responders (patients 11, 17, 29 and 30) had stable disease that was not confirmed on a subsequent scan.

This classification differs from two definitions used for the same trial. First, the trial reported a confirmed objective response rate of 9.4%[1], which requires a response to be confirmed on a second assessment; response categories based on the most recent assessment can therefore differ from confirmed best overall response. Second, Huffman et al.[1] defined clinical benefit as CR, PR, or SD lasting at least 6 months, under which the four patients with unconfirmed SD would be grouped with patients without clinical benefit. We used the most recent assessment so that the ctDNA analyses and the model analyses share one grouping, and we repeated the ctDNA comparisons under the clinical-benefit definition as a sensitivity analysis (Note S2).

**Note S2. ctDNA data handling and statistical comparisons**

Plasma TTMV-HPV DNA was measured with the NavDx assay (CareDx, Inc., Brisbane, CA) at baseline (C1D1), at the start of Cycle 2 (C2D1, Week 3), at the start of

Cycle 3 (C3D1, Week 6), and every other cycle thereafter. Results are reported as a TTMV-HPV DNA score, where 0 indicates that no TTMV-HPV DNA was detected; throughout, "ctDNA" refers to this score.

Three patients (11, 17 and 24) had undetectable ctDNA at every available timepoint, and one patient (27) had no ctDNA samples. These four patients were excluded from all ctDNA group comparisons, because a series that stays at zero carries no information about change and adds tied values at zero. Within each interval, patients without a sample at either timepoint were also excluded. This left 21 patients for the baseline analyses (10 responders, 11 non-responders), 20 for baseline to Cycle 2 (9, 11), 18 for baseline to Cycle 3 (9, 9), and 19 for Cycle 2 to Cycle 3 (10, 9). Patient-level values and the patients included in each comparison are given in Data File S1.

Early changes were quantified as the $\log_{10}$ fold change, $\log_{10}\left[\frac{(C_2+1)}{(C_1+1)}\right]$, where 1 is added to every score so that undetectable results can be included. Raw change and percent change were also computed and showed the same pattern of results (Data File S1). Group differences were tested with the two-sided Mann-Whitney U (Wilcoxon rank-sum) test using the normal approximation with continuity correction ($\alpha = 0.05$). The association between baseline ctDNA and baseline sum of longest diameters (SLD) was assessed with Spearman's rank correlation among patients with detectable baseline ctDNA ($n = 21$; $\rho = 0.67$).

Comparisons were made across three intervals and three change measures without correction for multiple testing, so these analyses should be regarded as exploratory.

**Note S3. Sensitivity, identifiability and fixed parameters**

Local sensitivity analysis[4] was used to rank the model parameters $\theta = \{\gamma_0, b, d, \delta, q, \varepsilon\}$ by their influence on the outputs $y = [V, C]$. The sensitivity matrix $S = \partial y/\partial\theta$ was computed numerically, and parameters were ranked by the time-averaged two-norm of their sensitivities. The tumor proliferation rate $b$ was the most sensitive parameter and the resistance-decay rate $\delta$ the least sensitive (Figure S2).

Identifiability was assessed with the correlation method[4]. From the covariance matrix $(S^{T}S)^{-1}$, pairwise correlation coefficients were computed, and a pair with an absolute correlation above 0.99 was considered non-identifiable, meaning the two parameters can compensate for each other without changing the fit. Two pairs exceeded this threshold: *b* and *d*, which affect tumor volume only through the net growth rate (*b* − *d*), and *q* and *ε*, which jointly set the ctDNA level relative to tumor volume (before treatment, ctDNA approaches $qdV/\varepsilon$ because clearance is fast).

One parameter from each pair was therefore fixed for all patients. The natural death rate *d* was fixed at the cohort mean from preliminary fits ($d$ = 0.064 day$^{-1}$). The clearance rate *ε* was fixed at 33 day$^{-1}$, which corresponds to a ctDNA half-life of $ln\ 2/\varepsilon \approx$ 30 minutes, consistent with reported ctDNA half-lives[2]. Because *d* is fixed, differences in *b* between patients should be read as differences in net tumor growth (*b* − *d*), not in proliferation alone. The remaining sensitive and identifiable parameters, $\gamma_0$, *b*, *δ* and *q*, were carried into the model-reduction step (Note S4).

**Note S4. Model reduction and selection**

Guided by the identifiability and sensitivity analyses, we held the natural tumor-cell death rate $d$ and the ctDNA elimination rate $\varepsilon$ fixed across all patients [$d$ fixed to the cohort-mean value, $\varepsilon$ fixed from[2]]. Beginning from a structure in which the four remaining identifiable parameters ($\gamma_0$, $b$, $\delta$, $q$) were patient-specific, we sequentially constrained one parameter at a time to a single uniform (cohort-wide) value and estimated the patient-specific values for the rest. Candidate structures were compared by normalized RMSE for both outputs and by AIC (Table S1).

With all four parameters patient-specific (**Structure 1**), the model achieved its closest agreement with the data ($nRMSE_V = 0.011$, $nRMSE_C = 0.030$) and, as expected for the least-constrained structure, the lowest AIC (123.10). Having the least coefficient of variation of CV = 1.08 [Supplementary Figure 3], the resistance-decay rate $\delta$ was set to a uniform value (**Structure 2**; patient-specific $\gamma_0$,$b$,$q$). This preserved good fits to both outputs ($nRMSE_V = 0.029$, $nRMSE_C = 0.039$). Further with a CV = 1.41 we constrained the initial treatment sensitivity $\gamma_0$ (**Structure 3**) and left only the tumor proliferation rate $b$

and the ctDNA shedding rate $q$ patient-specific [Supplementary Figure 4]. This structure reproduced both trajectories in individual patients, including representative responder and non-responder patients (**Figure 5a**), and across the cohort ($nRMSE_V = 0.042$, $nRMSE_C = 0.078$), with simulated and measured values clustering about the line of identity for both volume and ctDNA (**Figure 5b**). As a sanity check on model behavior, we also simulated each patient without treatment (dashed curves); as expected, volume and ctDNA grew exponentially in the absence of treatment. Fits for all patients are shown in [Supplementary Figure 2].

As parameters were moved from patient-specific to uniform, AIC increased across the series $(123.10 - 172.83 - 217.83)$ while fit error rose only modestly, reflecting the expected trade-off between goodness of fit and model complexity: the fullest model fits best, whereas the reduced models describe the cohort with substantially fewer free parameters. Given this, we selected **Structure 3** as the final parameter structure [Supplementary Figure 5]. Although it does not minimize AIC, it captures the full cohort with only two patient-specific parameters – reducing the degrees of freedom substantially while retaining good accuracy across all patients – and isolates the biological differences between patients to tumor proliferation $(b)$ and ctDNA shedding $(q)$, with all treatment-sensitivity parameters shared cohort-wide.

It is important to note that across all three structures, the tumor proliferation rate $b$ was significantly higher in non-responders than in responders. The final optimization of Structure 3 showed that the ctDNA shedding rate $q$ did not significantly differ between patients **(Figure 5c)**.

The difference in *b* did not depend on which parameters were shared: *b* was significantly higher in non-responders in Structure 1 ($p = 0.010$), Structure 2 ($p = 0.010$) and Structure 3 ($p = 0.003$) (Figures S3–S5). In Structure 3, the median *b* was 0.070 day$^{-1}$ in responders and 0.102 day$^{-1}$ in non-responders (n = 10 and 11 patients with a usable ctDNA series). With *d* fixed at 0.064 day$^{-1}$, this corresponds to untreated tumor-volume doubling times of roughly 120 days in responders and 18 days in non-responders. The ctDNA shedding rate *q* varied over several orders of magnitude between patients (CV = 3.56) and did not differ between groups (median $9.3 \times 103 \text{ vs.} 2.3 \times 104$; $p =$

$0.19$). Given the small cohort, a non-significant result for $q$ should be read as inconclusive rather than as evidence of no difference.

**Figure S1**

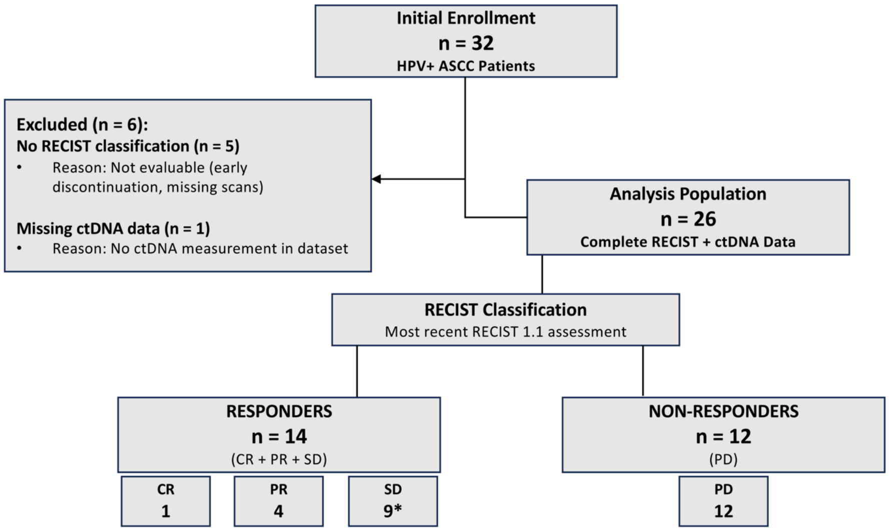


**Figure S1. CONSORT diagram**. Of 32 enrolled patients from the NCT02919969 trial[1], 26 had both RECIST and ctDNA data. Patients were classified by their most recent RECIST 1.1 assessment as responders (CR, PR or SD; n = 14, including 4 with unconfirmed SD) or non-responders (PD; n = 12).

**Figure S2: Sensitivity Analysis**

Local sensitivity analysis ranked the tumor proliferation rate $b$ as the most sensitive parameter and the resistance-decay rate $\delta$ as the least sensitive. Correlation analysis found correlations between *b* and *d*, which affect tumor volume only through their net difference (b − d) and with $\varepsilon$ and $q$, which affect ctDNA shedding and clearance. After accounting for these, the sensitive and identifiable parameters were $[\gamma_0,\ b,\ \delta,\ q]$.

**Figure S3: Model Structure 1**

With all four parameters patient-specific, Structure 1, the model achieved its closest agreement with the data ($nRMSE_V = 0.011$, $nRMSE_C = 0.030$) and, as expected for the least-constrained structure, the lowest AIC (123.10).

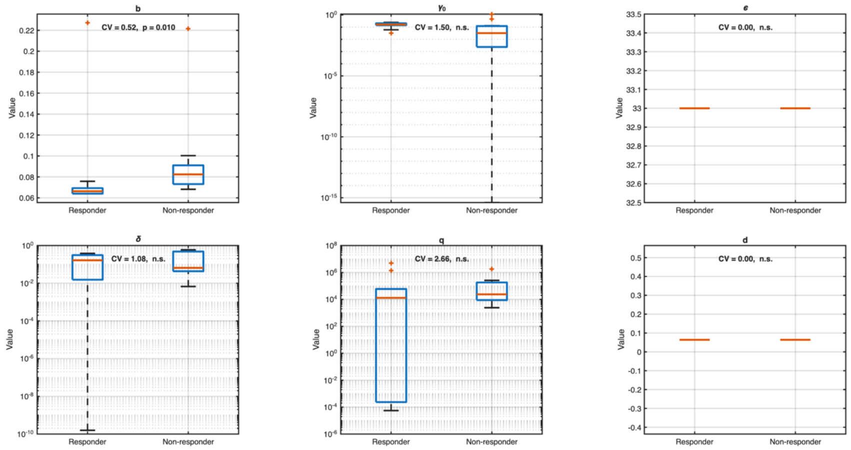


**Figure S3. Model 1. Parameter distributions by response group.** Box plots of patient-specific parameters ($b, \gamma_0, \delta, q$) for responders and non-responders. $d$ (0.0642 day⁻¹) and $\varepsilon$ (33 day⁻¹) were fixed. Each panel shows the coefficient of variation (CV) across patients and the between-group p-value. Only the proliferation rate b differed significantly between groups (p = 0.010).

**Figure S4: Model Structure 2**

From Structure 1, with the resistance-decay rate $\delta$ having the least coefficient of variation of CV = 1.08, we set that parameter to be a uniform value thus having (patient-specific $\gamma_0$ ,$b$,$q$). This preserved good fits to both outputs ($nRMSE_V = 0.029$, $nRMSE_C = 0.039$).

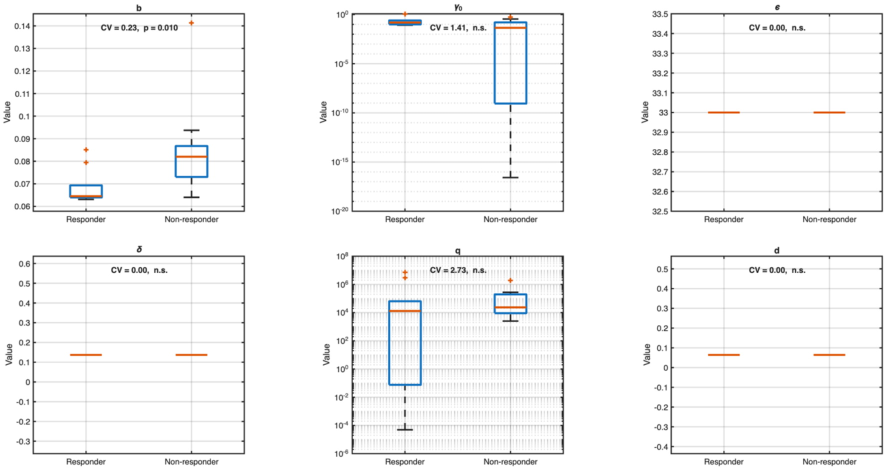


**Figure S4. Model 2 parameter distributions by response group.** Box plots of the patient-specific parameters ($b, \gamma_0, q$) for responders and non-responders. $\delta$ was held uniform across patients at the cohort value from Model 1. $d$ (0.0642 day⁻¹) and $\varepsilon$ (33 day⁻¹) were fixed. Each panel shows the coefficient of variation (CV) across patients and the between-group p-value. Only the proliferation rate $b$ differed significantly between groups (p = 0.010).

**Figure S5: Model Structure 3**

Further from structure 2, with a CV = 1.41 we constrained the initial treatment sensitivity $\gamma_0$ and left only the tumor proliferation rate $b$ and the ctDNA shedding rate $q$ patient-specific. This structure reproduced both trajectories in individual patients, including representative responder and non-responder patients, and across the cohort ($nRMSE_V = 0.042$, $nRMSE_C = 0.078$), with simulated and measured values clustering about the line of identity for both volume and ctDNA.

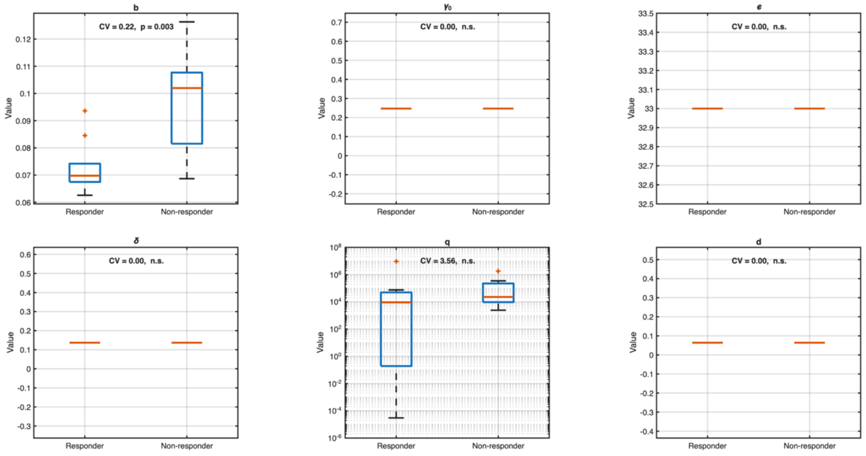


**Figure S5. Model 3 parameter distributions by response group.** Box plots of the patient-specific parameters $(b, q)$ for responders and non-responders. $\gamma_0$ and $\delta$ were held uniform across patients at the cohort values from Model 2. $d$ $(0.0642$ day⁻¹$)$ and $\varepsilon$ $(33$ day⁻¹$)$ were fixed. Each panel shows the coefficient of variation (CV) across patients and the between-group p-value. Only the proliferation rate $b$ differed significantly between groups ($p = 0.003$).

Table S1:

| Model | Uniform | Optimized (PS) | nRMSE(V) | nRMSE(C) | AIC |
|---|---|---|---|---|---|
| 1 | - | $\gamma_0$, b, $\delta$, q | 0.011 | 0.030 | 123.1023 |
| 2 | $\delta$ | $\gamma_0$, b, q | 0.029 | 0.039 | 172.8318 |
| 3 | $\delta, \gamma_0$ | b, q | 0.042 | 0.078 | 217.8334 |

**Table S1. Model Structure comparison**. For each model candidate structure, we have the parameters fixed to a uniform (cohort-wide) value, the patient-specific (optimized) parameters, normalized RMSE for volume and ctDNA, AIC, and whether responder/non-responder parameter differences were significant.

**Commented [C1]:** The caption says the table shows "whether responder/non-responder parameter differences were significant", but the table has no such column. Either add a column (b: p = 0.010, 0.010, 0.003; all other parameters n.s.) or remove that phrase.

**Data File S1 (separate Excel file)**

Data File S1 contains the patient-level ctDNA data and all group comparisons reported in the main text and in Note S2. Its sheets are:

File 1 – Patient groupings, exclusions, and the number of patients in each ctDNA comparison, with the clinical-benefit grouping used for the sensitivity analysis.

File 2 – Patient-level TTMV-HPV DNA scores at C1D1, C2D1 and C3D1, with raw change, $\log_{10}$ fold change and percent change for each interval.

File 3 – Mann-Whitney U calculations (ranks, U, z and p-values), medians, and the number of patients with a rising score, for responders versus non-responders.

File 3b – The same comparisons under the clinical-benefit grouping (sensitivity analysis).

Files 4 and 5 – Summary tables of the comparisons in File 3.

File 6 and S3–S5 – Model parameter comparisons between responders and non-responders for Structures 1–3.

**References**


1 Huffman, B. M. *et al.* Biomarkers of pembrolizumab efficacy in advanced anal squamous cell carcinoma: analysis of a phase II clinical trial and a cohort of long-term responders. *J Immunother Cancer* **12** (2024). https://doi.org/10.1136/jitc-2023-008436

2 Li, A., Lou, E., Leder, K. & Foo, J. Early Circulating Tumor DNA Kinetics as a Dynamic Biomarker of Cancer Treatment Response. *JCO Clin Cancer Inform* **9**, e2400160 (2025). https://doi.org/10.1200/cci-24-00160

3 Eisenhauer, E. A. *et al.* New response evaluation criteria in solid tumours: revised RECIST guideline (version 1.1). *Eur J Cancer* 45, 228-247 (2009). https://doi.org/10.1016/j.ejca.2008.10.026

4 Olufsen, M. S. & Ottesen, J. T. A practical approach to parameter estimation applied to model predicting heart rate regulation. *J Math Biol* 67, 39-68 (2013). https://doi.org/10.1007/s00285-012-0535-8
